# Development of fluoroquinolone resistance through antibiotic tolerance in *Campylobacter jejuni*

**DOI:** 10.1101/2022.05.21.492921

**Authors:** Myungseo Park, Jinshil Kim, Jill Feinstein, Kevin S. Lang, Sangryeol Ryu, Byeonghwa Jeon

## Abstract

Antibiotic tolerance not only enables bacteria to survive under acute antibiotic exposures but also provides bacteria with a window of time to develop antibiotic resistance. *Campylobacter jejuni* is increasingly resistant to clinically important antibiotics, particularly fluoroquinolones (FQs). Currently, little is known about antibiotic tolerance and its effects on resistance development in *C. jejuni*. Here, we show that exposure to ciprofloxacin and tetracycline at concentrations 10 and 100 times higher than the minimum inhibitory concentration (MIC) induce antibiotic tolerance in *C. jejuni*, whereas gentamicin and erythromycin treatment cause cell death. Interestingly, FQ resistance is rapidly developed in *C. jejuni* after tolerance induction by ciprofloxacin and tetracycline. Furthermore, alkyl hydroperoxide reductase plays a critical role in preventing FQ resistance development in *C. jejuni* during antibiotic tolerance by alleviating oxidative stress. Together, these results demonstrate that exposure of *C. jejuni* to antibiotics used to treat campylobacteriosis can induce antibiotic tolerance and that FQ-resistant (FQ^R^) *C. jejuni* rapidly emerges through tolerance induction by FQs and non-FQ antibiotics. Work presented in this study shows mechanisms underlying the high prevalence of FQ^R^ *C. jejuni* and provides an insight into the effects of antibiotic tolerance on resistance development.

**Importance:** Antibiotic tolerance compromises the efficacy of antibiotic treatment by extending bacterial survival and developing mutations associated with antibiotic resistance. Despite growing public health concerns about antibiotic resistance in *C. jejuni*, antibiotic tolerance has not yet been investigated in this important zoonotic pathogen. Here, our results show that exposure of *C. jejuni* to ciprofloxacin and tetracycline, a common agricultural antibiotic, develops antibiotic tolerance, which subsequently facilitates the emergence of FQ^R^ *C. jejuni*. Since antibiotic-resistant *C. jejuni* is transmitted primarily from animals to humans, our study suggests that non-FQ drugs, such as tetracycline, used for animals can also promote FQ resistance development by inducing antibiotic tolerance in *C. jejuni*. Overall, the findings in this study help us understand mechanisms of resistance development through the induction of antibiotic tolerance.

## INTRODUCTION

Antibiotic tolerance enables antibiotic-susceptible bacteria to survive acute exposure to high doses of bactericidal antibiotics (1). Whereas tolerance refers to the ability of an entire bacterial population to withstand antibiotic treatment for prolonged periods of time, persistence indicates tolerance in a subpopulation of a clonal bacterial population (2). Extended survival of pathogenic bacteria under antibiotic treatment, whether by tolerance or persistence, can cause prolonged and recurrent infections, resulting in adverse clinical outcomes and treatment failure (1, 2).

Furthermore, the extended survival of tolerant and persistent bacteria under antibiotic treatment can provide a window of time for the emergence of antibiotic-resistant bacteria (3, 4).

*Campylobacter* spp., particularly *Campylobacter jejuni*, is a leading bacterial cause of gastroenteritis and accounts for 400-500 million cases of diarrhea worldwide per year (5). *C. jejuni* colonizes the gastrointestinal tracts of a wide range of livestock and companion animals, both mammals and avian species, and is transmitted to humans mainly through foodborne routes (6). In addition, the increasing prevalence of *Campylobacter* isolates resistant to clinically important antibiotics is another serious public health concern. Importantly, the prevalence of fluoroquinolone-resistant (FQ^R^) *Campylobacter* increases at an alarming rate and significantly compromised the efficacy of this critically important antibiotic class (7). Approximately 28.5% of 1.5 million campylobacteriosis cases in the United States are associated with FQ resistance (8), and FQ^R^ *Campylobacter* causes adverse patient outcomes (9, 10). In some countries, FQ^R^ *Campylobacter* is already widespread, such as 76% in Italy (11), 87% in China (12), and 89% in Thailand (13). The World Health Organization (WHO) classified FQ^R^ *Campylobacter* as one of the high-priority pathogens for which new antimicrobials should be developed (14).

Antibiotic tolerance not only extends the survival of bacteria under antibiotic treatment but also contributes to the emergence of antibiotic-resistant bacteria (3, 4). Despite public health concerns about antibiotic resistance in *Campylobacter*, little is known about whether *Campylobacter* can develop antibiotic tolerance and, if it does, how tolerance affects the development of antibiotic resistance in this highly mutable bacterium. In this study, we demonstrate that *C. jejuni* develops antibiotic tolerance by exposure to high concentrations of antibiotics. Importantly, FQ resistance is rapidly developed through antibiotic tolerance induced by FQs and non-FQ antibiotics.

## RESULTS

### Tolerance induction in *C. jejuni* by exposure to high concentrations of antibiotics

We wanted to investigate if *C. jejuni* could develop tolerance by antibiotic exposure. FQs, erythromycin, tetracycline, and gentamicin were selected for treatment because FQs are the most commonly used oral antibiotic for empirical treatment of gastroenteritis (15), erythromycin is the drug of choice for treating campylobacterosis, and gentamycin and tetracyclines are alternative drugs to treat systemic infections with *Campylobacter* (16). To this end, we treated *C. jejuni* cultures with these antibiotics at concentrations 10 and 100 times higher than the minimum inhibitory concentration (MIC) and measured the viability of *C. jejuni*. We found that *C. jejuni* developed tolerance to ciprofloxacin, a FQ drug, and tetracycline but was killed by gentamicin and erythromycin (Fig. 1). Translation inhibitors, such as tetracycline and erythromycin, are generally considered bacteriostatic but exhibited bactericidal activity when used at high concentrations (Fig. 1B, C). None of the antibiotics used did not produce biphasic killing curves (Fig. 1), which are unique to antibiotic persistence due to the presence of subpopulations. This indicates that antibiotic treatment induced tolerance, not persistence, in *C. jejuni*. Interestingly, treatment with high concentrations (i.e., 10x and 100x MICs) of ciprofloxacin did not lead to immediate bacterial killing and gradually inhibited *C. jejuni* over time but increased the level of *C. jejuni* after 48 hours due to the emergence of FQ^R^ *C. jejuni* (Fig. 1A). These results show that tolerance can be induced in *C. jejuni* by exposure to high concentrations of ciprofloxacin and tetracycline, and FQ resistance rapidly develops during antibiotic tolerance.

**Fig. 1.**
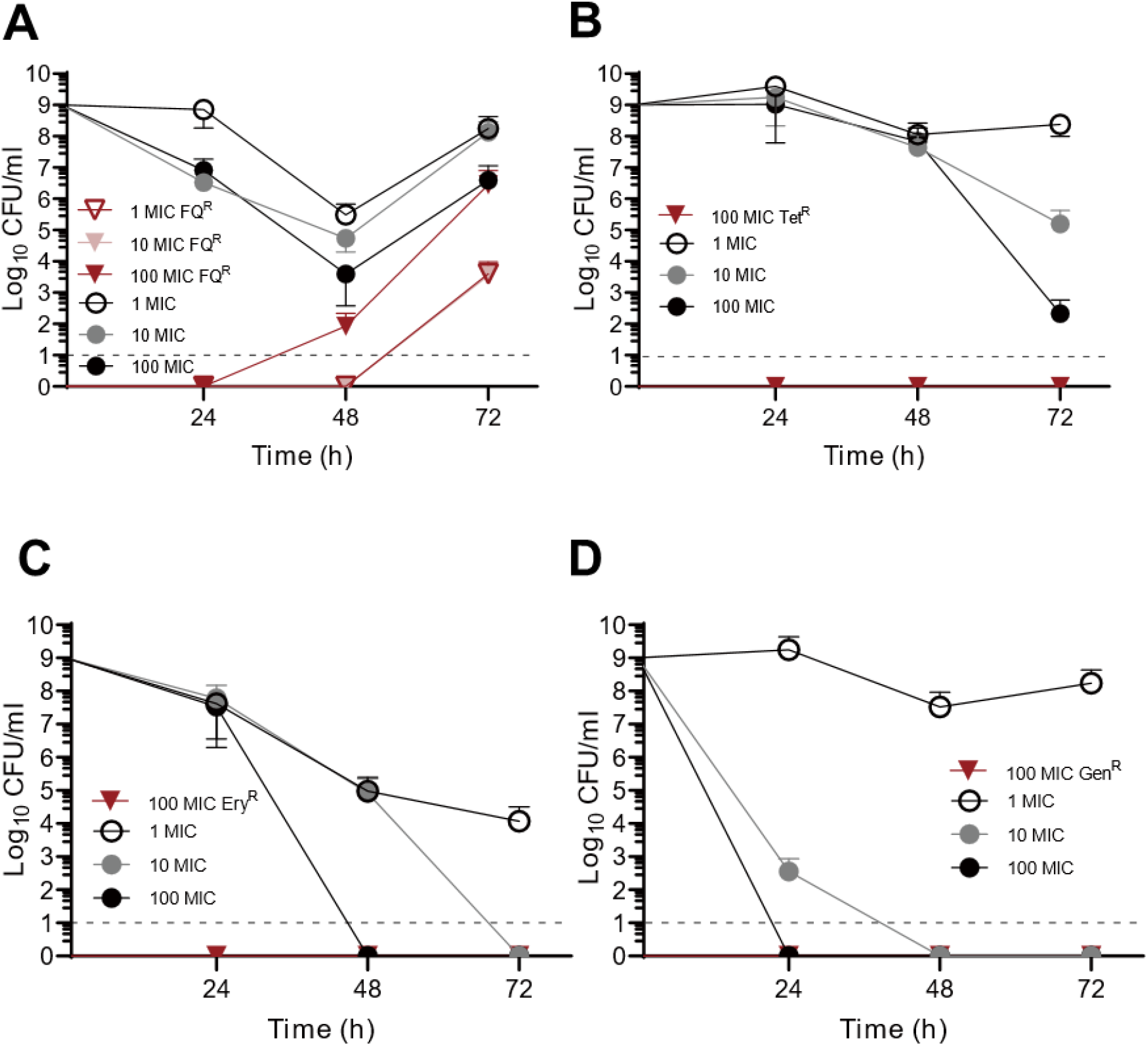
Induction of antibiotic tolerance in *C. jejuni* by exposure to high concentrations of (A) ciprofloxacin, (B) tetracycline, (C) erythromycin, and (D) gentamicin. The results show the means and standard deviations of the results of three independent experiments. Black lines indicate the total *C. jejuni* levels, and red lines show the levels of *C. jejuni* resistant to the antibiotic used for the treatment. Antibiotic concentrations were calculated based on the MICs of ciprofloxacin, tetracycline, erythromycin, gentamicin in *C. jejuni* NCTC 11168, which were 0.063 μg/ml, 0.031 μg/ml, 1 μg/ml, and 0.5 μg/ml, respectively. Dotted lines are the detecting limit.

### Hydroxyl radical level decides between tolerance induction and bacterial killing in *C. jejuni*

Since hydroxyl radical formation is a general mechanism for bacterial lethality by bactericidal antibiotics (17), we hypothesized that *C. jejuni* must overcome increased oxidative stress after acute antibiotic exposure to maintain tolerance, or the level of hydroxy radicals should not exceed a lethal threshold during tolerance. To test the hypothesis, we measured hydroxyl radicals in *C. jejuni* during exposure to high levels of antibiotics. As predicted, the level of hydroxyl radicals was significantly elevated by exposure to 10x MIC of erythromycin and gentamicin (Fig. 2), which accounts for rapid killing of *C. jejuni* by these antibiotics (Fig. 1C, D). Compared to these antibiotics, ciprofloxacin and tetracycline induced the formation of hydroxyl radicals at lower levels (Fig. 2). These results show that the level of hydroxyl radical formation correlates with whether antibiotic treatment leads to bacterial killing or induces antibiotic tolerance in *C. jejuni*.

**Fig. 2.**
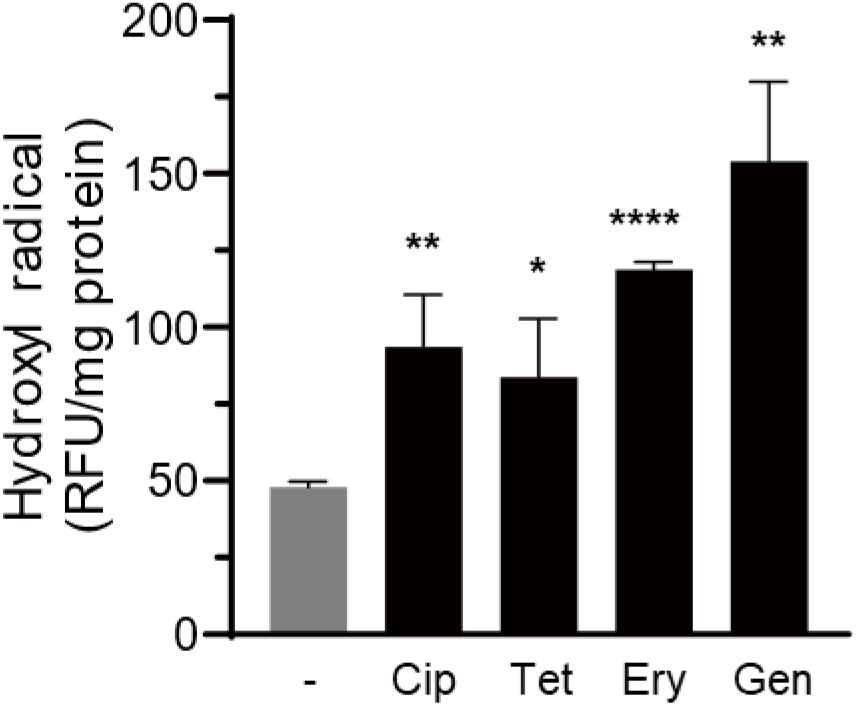
Hydroxyl radical formation during antibiotic exposure in *C. jejuni*. The level of hydroxyl radicals was measured after exposure to 10x MICs of ciprofloxacin (Cip; 0.63 μg/ml), tetracycline (Tet; 0.31 μg/ml), erythromycin (Ery; 10 μg/ml), and gentamicin (Gen; 5 μg/ml) for 24 hours. The results show the means and standard deviations of an experiment with three samples. The experiments were repeated three times and produced similar results. Statistical analysis was conducted with Student’s *t*-test in comparison with a non-treated control. *: *P*<0.05, **: *P*<0.01, and ****: *P*<0.0001.

### FQ resistance development by exposure to tolerance-inducing antibiotics in *C. jejuni*

Oxidative stress during antibiotic treatment can not only affect bacterial killing but also cause DNA mutations (17). Since ciprofloxacin treatment resulted in the emergence of FQ^R^ *C. jejuni* (Fig. 1A) and escalated oxidative stress in *C. jejuni* (Fig. 2), we speculated that oxidative stress increase by exposure to tolerance-inducing antibiotics developed DNA mutations conferring FQ resistance. We also hypothesized that *C. jejuni* might develop FQ resistance by exposure to non-FQ drugs because treatment with non-FQ drugs leads to hydroxyl radical formation (Fig. 2). To test this hypothesis, we exposed *C. jejuni* to 100x MIC ciprofloxacin, 100x MIC tetracycline, 100x MIC erythromycin, and 10x MIC gentamicin; we decreased the gentamicin concentration because of its strong bactericidal activity (Fig. 1D). Interestingly, FQ^R^ *C. jejuni* clones rapidly emerged after tolerance induction by ciprofloxacin and non-FQ drugs (Fig. 3A). Remarkably, the level of FQ^R^ cells arising during tetracycline treatment was significantly higher than that in a control without antibiotic treatment (Fig. 3A). The population ratio of FQ^R^ *C. jejuni* to total *C. jejuni* was significantly increased in the presence of ciprofloxacin over 48 hours (Fig. 3B). This suggests that FQ^R^ *C. jejuni* populations are enriched in the presence of ciprofloxacin during antibiotic tolerance. The emergence of FQ^R^ *C. jejuni* was limitedly observed in the presence of antibiotics with strong bactericidal activity. Whereas gentamycin treatment rapidly killed *C. jejuni*, erythromycin induced the emergence of FQ^R^ *C. jejuni* clone sporadically (one in three experiments) after 24 hours, but FQ^R^ *C. jejuni* was not detected after 48 hours owing to bacterial killing by erythromycin (Fig. 3B). Altogether, these results suggest that FQ resistance is rapidly developed in *C. jejuni* during antibiotic tolerance induced by FQs and non-FQ antibiotics.

**Fig. 3.**
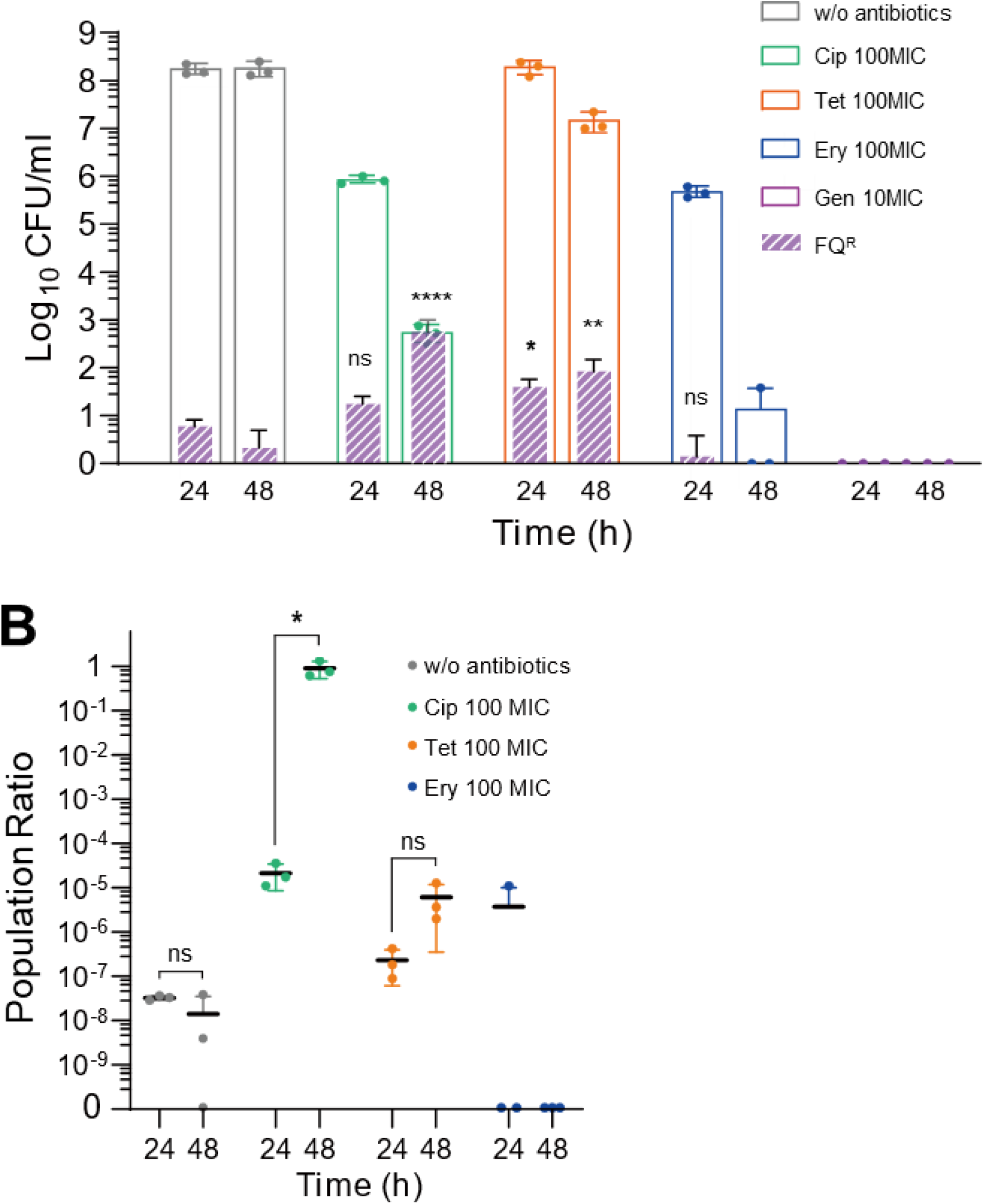
Development of fluoroquinolone (FQ) resistance during antibiotic tolerance in *C. jejuni*. (**A**) The emergence of FQ-resistant (FQ^R^) *C. jejuni* under treatment with 100x MICs of ciprofloxacin (Cip; 6.3 μg/ml), tetracycline (Tet; 3.1 μg/ml), and erythromycin (Ery; 100 μg/ml), and 10x MIC of gentamicin (Gen; 5 μg/ml). Gen was treated at a lower concentration due to its strong bactericidal activity. The results show the means and standard deviations of the levels of total *C. jejuni* (empty bars) and FQ^R^ *C. jejuni* (patterned fills) of the results from three independent experiments. Statistical analysis was performed with Student’s *t*-test in comparison with a non-treated control at the same sampling time. ns: non-significant, *: *P*<0.05, **: *P*<0.01, and ****: *P*<0.0001. (**B**) Enrichment of FQ^R^ *C. jejuni* during antibiotic tolerance. The results show the ratio of FQ^R^ *C. jejuni* to the total *C. jejuni*. Statistical analysis was conducted with Student’s *t*-test. ns: non-significant, *: *P*<0.05.

### AhpC reduces FQ resistance development in *C. jejuni* during antibiotic tolerance

Our results suggest that augmented oxidative stress during antibiotic exposure can be responsible for the emergence of FQ^R^ *C. jejuni* in antibiotic-tolerant cells. *C. jejuni* harbors a sole copy of genes encoding alkyl hydroperoxide reductase (AhpC), catalase (KatA), and superoxide dismutase (SodB)(18); all involved in the detoxification of different reactive oxygen species. Using *ahpC, katA*, and *sodB* knockout mutants, we examined which antioxidant enzyme plays a major role in the prevention of FQ resistance development in *C. jejuni* during antibiotic tolerance.

Remarkably, an *ΔahpC* mutation significantly increased the frequency of FQ resistance development compared to wild type (WT; Fig. 4A), whereas *ΔkatA* and *ΔsodB* mutations did not affect the development of FQ resistance compared to WT (Fig. S1). In the *ΔahpC* mutant, ciprofloxacin treatment markedly increased the accumulation of hydrogen peroxide (Fig. 4B), a substrate of AhpC, and hydroxyl radicals (Fig. 4C). These results suggest that increased oxidative stress facilitates the development of FQ resistance during antibiotic tolerance in *C. jejuni* and AhpC plays an important role in reducing the development of FQ resistance by alleviating oxidative stress.

**Fig. 4.**
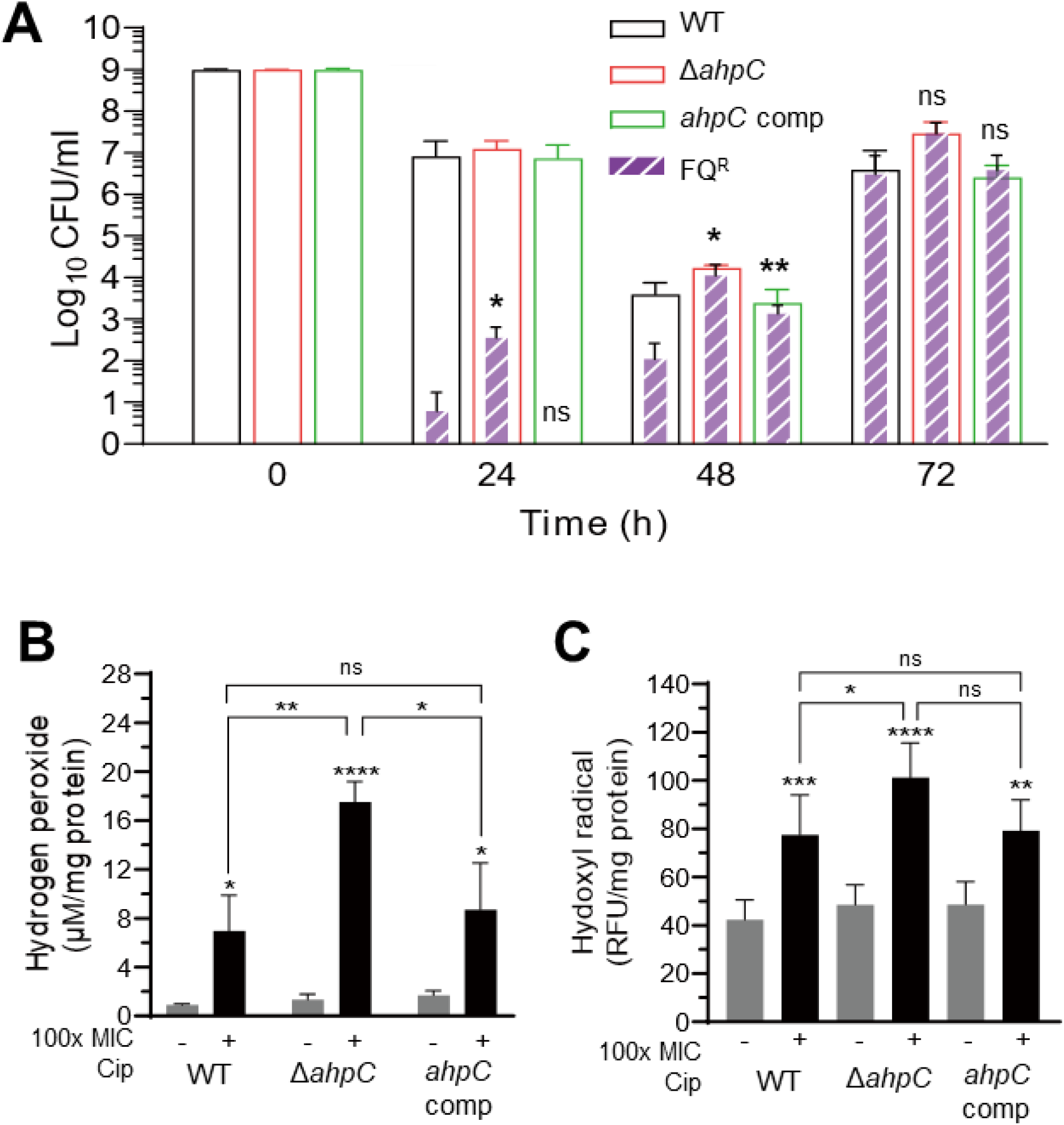
Enhanced development of fluoroquinolone (FQ) resistance in an *ΔahpC* mutant during antibiotic tolerance induced by ciprofloxacin. (**A**) Significant increase in FQ resistance development in an *ΔahpC* mutant during antibiotic tolerance induced by 100x MIC of ciprofloxacin (6.3 μg/ml). The results show the means and standard deviations of the levels of total *C. jejuni* (empty bars) and FQ^R^ *C. jejuni* (patterned fills) of the results from three independent experiments. Statistical analysis was conducted with Student’s *t*-test in comparison with WT. ns: non-significant, *: *P*<0.05, and **: *P*<0.01. *ahpC* comp: *ahpC*-complemented strain. Increased production of hydrogen peroxide (**B**) and hydroxyl radicals (**C**) in an Δ*ahpC* mutant. The + and – signs indicate the presence and absence of 100x MIC of ciprofloxacin (6.3 μg/ml), respectively. Statistical analysis was conducted with Student’s *t*-test. ns: non-significant, *: *P*<0.05, **: *P*<0.01, ***: *P*<0.001, and ****: *P*<0.0001.

### Oxidative stress defense regulators modulating *ahpC* transcription affect FQ resistance development during antibiotic tolerance in *C. jejuni*

To further confirm the role of AhpC in FQ resistance development during antibiotic tolerance in *C. jejuni*, we used mutants defective in the regulation of oxidative stress defense. *C. jejuni* lacks OxyR and SoxRS, the most common regulators of oxidative stress defense in many Gram-negative bacteria (19, 20), and uses PerR (21) and CosR (22) to respond to oxidative stress. PerR is a repressor of *ahpC* transcription (21, 23). CosR is a response regulator positively regulating *ahpC* transcription (22). Thus, a *ΔperR* mutation increases *ahpC* transcription by derepression, and CosR-overexpression increases *ahpC* transcription by positive regulation. We measured the development of FQ resistance after inducing antibiotic tolerance in a CosR-overexpression strain (Fig. 5A) and a *ΔperR* mutant (Fig. 5B). The frequency of FQ resistance development was substantially reduced in the *ΔperR* and CosR-overexpression mutants. However, when *ahpC* was deleted in these mutants, the frequency of FQ resistance was increased to levels similar to that of WT (Fig. 5), confirming the role of AhpC in the control of FQ resistance development during antibiotic tolerance. Moreover, we observed the level of *ahpC* transcription was significantly increased in antibiotic-tolerant *C. jejuni* under treatment with high concentrations of ciprofloxacin, tetracycline, erythromycin, and gentamicin (Fig. S2). This indicates the antioxidation function of AhpC is required during antibiotic tolerance in *C. jejuni*. These results demonstrate that AhpC plays an important role in preventing the development of FQ resistance through tolerance induction in *C. jejuni*.

**Fig. 5.**
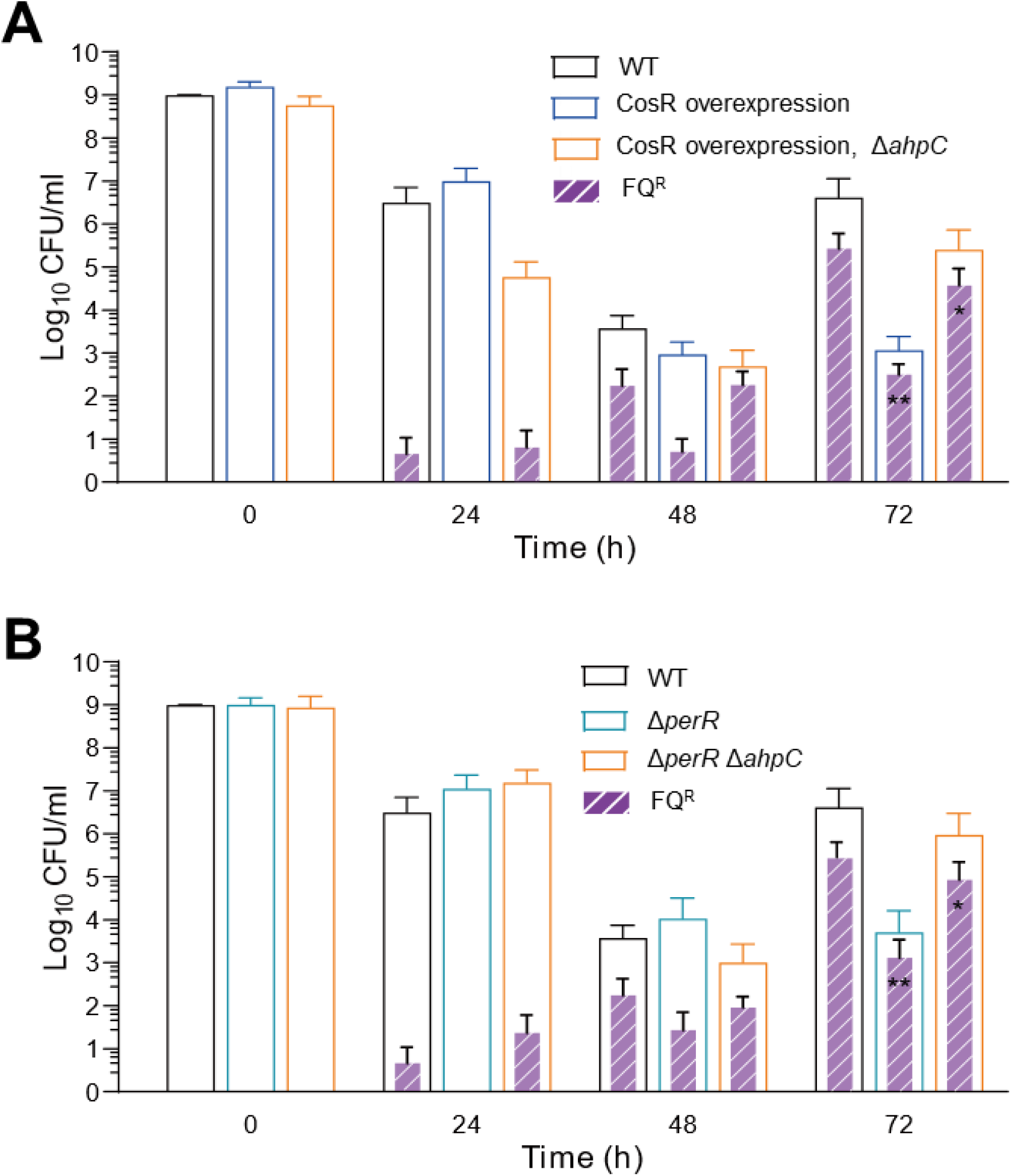
Effects of *ahpC* regulation on FQ resistance development in *C. jejuni* during antibiotic tolerance induced by ciprofloxacin. (**A**) FQ resistance development in a CosR-overexpression strain and a CosR-overexpression strain with an Δ*ahpC* mutation. (**B**) FQ resistance development in a Δ*perR* mutant and a Δ*perR* Δ*ahpC* mutant. The concentration of ciprofloxacin is 100x MIC (6.3 μg/ml). The results show the means and standard deviations of the levels of total *C. jejuni* (empty bars) and FQ^R^ *C. jejuni* (patterned fills) of the results from three independent experiments. Statistical analysis was conducted with two-way ANOVA followed by Dunnett’s multiple comparisons test. *: *P*<0.05, **: *P*<0.01.

## DISCUSSION

The data in this study show that antibiotic tolerance induced by exposure to FQs and non-FQ drugs promotes the development of FQ resistance in *C. jejuni*. Studies demonstrate that antibiotic tolerance or persistence induced by intermittent antibiotic exposures or drug combinations precedes the emergence of antibiotic resistance mutations in *Escherichia coli* and *Staphylococcus aureus* (3, 4). In *E. coli*, FQs induce the SOS responses in persisters and increase DNA mutations through the induction of error-prone DNA polymerase V (24). However, *C. jejuni* lacks SOS response systems, and error-prone DNA polymerases have not been reported in *C. jejuni*. Instead, our data exhibit that AhpC plays a critical role in preventing FQ resistance development. AhpC is involved in the detoxification of organic peroxides and low physiological levels of hydrogen peroxide (25). Deletion of *ahpC* results in the accumulation of hydrogen peroxide and hydroxyl radicals (Fig. 4B and C), which increases the frequency of FQ resistance development during antibiotic tolerance in *C. jejuni* (Fig. 4A). Hydrogen peroxide is generated by accidental autooxidation of non-respiratory flavoproteins (26, 27) and dismutation of superoxide (28), and is converted to hydroxyl radicals through the iron-catalyzed Fenton reaction (28). Thus, enzymatic degradation of hydrogen peroxide by AhpC can reduce oxidative DNA damage by toxic hydroxyl radicals and consequently decreases FQ resistance development during antibiotic tolerance in *C. jejuni* (Fig. 6).

**Fig. 6.**
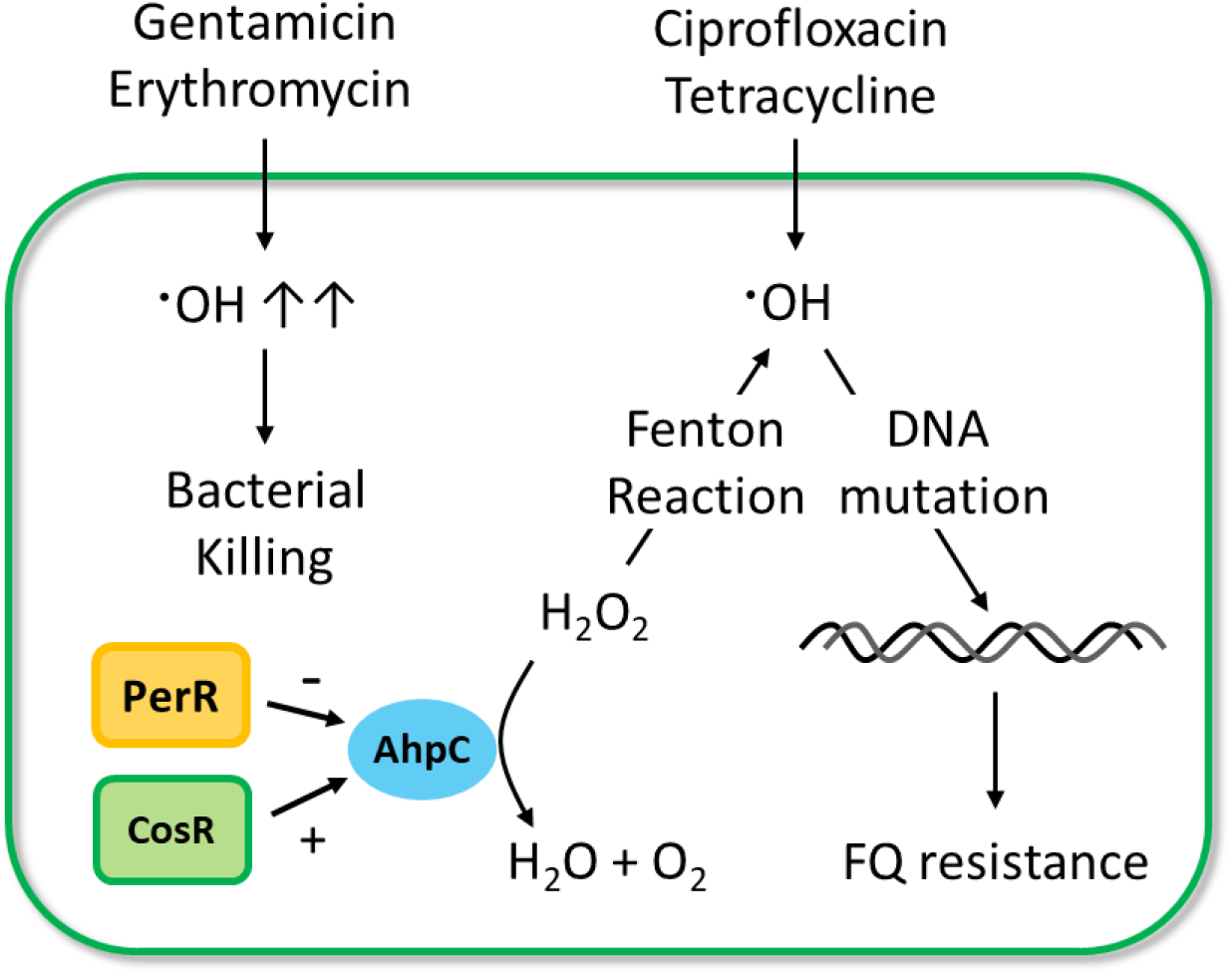
Schematic diagram of FQ resistance development through tolerance induction in *C. jejuni*. Exposure to high concentrations of gentamycin and erythromycin induces the formation of hydroxyl radicals at high levels, which lead to cell death. Ciprofloxacin and tetracycline exposures do not generate lethal levels of hydroxyl radicals and cause DNA mutations conferring FQ resistance. AhpC decreases the level of hydroxyl radicals by degrading hydrogen peroxide, which can be converted to hydroxyl radicals by the Fenton reaction. The *ahpC* transcription is negatively and positively regulated by PerR and CosR, respectively.

FQ^R^ *C. jejuni* rapidly emerged after tolerance induction, but the size of FQ^R^ *C. jejuni* population altered differentially over time depending on the antibiotic used for tolerance induction (Fig. 3). FQ^R^ *C. jejuni* appeared within 24 hours under tetracycline treatment and remained at a similar ratio until 48 hours (Fig. 3B), whereas the population of FQ^R^ *C. jejuni* was significantly increased over time in the presence of ciprofloxacin (Fig. 3B). This can be ascribed to selective pressure made by ciprofloxacin. Ciprofloxacin treatment killed FQ-susceptible *C. jejuni* over time, increasing the ratio of FQ^R^ *C. jejuni* to the total *C. jejuni* (Fig. 3B). Due to growth arrest in tolerant cells under antibiotic treatment, the increase in FQ^R^ *C. jejuni* cells can be driven by further oxidative damage by extended antibiotic exposure. Additionally, selective enrichment appears to play a role in increasing the number of FQ^R^ *C. jejuni* because FQ^R^ *C. jejuni* populations did not increase significantly under tetracycline treatment (Fig. 3B). These data suggest *C. jejuni* may not be in complete dormancy during antibiotic tolerance. In this regard, further studies are needed to understand the physiology of antibiotic-tolerant *C. jejuni*.

FQs are the most commonly used oral antibiotic to treat various bacterial infections (15, 29). FQs bind to DNA gyrase and topoisomerase IV, generate double-stranded breaks in DNA, and lead to bacterial cell death (30, 31); thus, DNA mutations leading to structural changes in topoisomerase IV and DNA gyrase are the major cause of FQ resistance (32, 33). However, genes encoding topoisomerase IV are absent in *C. jejuni* (18, 34), and mutations in DNA gyrase subunit B do not mediate FQ resistance in *C. jejuni* (35). Instead, *C. jejuni* develops FQ resistance by single-step point mutations in GyrA, DNA gyrase subunit A (35, 36). FQ treatment of chickens colonized by *C. jejuni* cannot completely eliminate *C. jejuni*, Instead, FQ^R^ *C. jejuni* emerges within a few days after FQ treatment and increases the level of *C. jejuni* (36). This is the same pattern of bacterial killing and FQ resistance development through antibiotic tolerance in our study (Fig. 1A), suggesting antibiotic tolerance may contribute to FQ resistance development *in vivo*.

The high prevalence of FQ^R^ *Campylobacter* has been ascribed mainly to the use of FQs in agriculture because *C. jejuni* is mainly transmitted from food-producing animals to humans. Considerable efforts have been made to curb FQ resistance in *Campylobacter* by banning FQs in poultry production and agriculture sectors in the United States and other countries. However, FQ resistance in *Campylobacter* remains increasing (37). This has been explained by the fitness benefits of FQ resistance in *C. jejuni*. Once *C. jejuni* develops FQ resistance, FQ^R^ *C. jejuni* outcompetes FQ-sensitive *C. jejuni* in the intestines of chickens, the primary reservoir for *C. jejuni*, even in the absence of FQs (38). This makes FQ^R^ *C. jejuni* prevalent in chickens. In addition, our results show that the induction of antibiotic tolerance is an important mechanism for the rapid development of FQ resistance in *C. jejuni*.

Notably, antibiotic tolerance induced by non-FQ drugs, particularly tetracycline, can facilitate the emergence of FQ^R^ *C. jejuni*. Tetracyclines are most widely used in agriculture, accounting for 66% and 31% of marketed agricultural antimicrobials in the United States (39) and the European Union (40), respectively. In the United States, the use of tetracyclines as growth promoters is no longer allowed and can be used only for therapeutic purposes. However, tetracyclines represent the largest volume of domestic sales of medically important antibiotics approved for use in food-producing animals, and about 3,948 tons of tetracyclines are sold and distributed for veterinary purposes in the country in 2020 (39). After medication in feed, more than 70 % of tetracyclines are unmetabolized and excreted from animals into the environment (41). This increases the chances of bacterial exposure to high levels of tetracyclines both inside and outside of animals. Many countries widely use tetracyclines as in-feed antibiotics for growth promotion, disease prevention, and treatment (42). Although FQs have been banned in livestock production in many countries, the widespread use of tetracycline or other tolerance-inducing antibiotics in animals can contribute to the development of FQ resistance in *C. jejuni*.

In summary, work in our study suggests that tetracyclines, an antibiotic class commonly used in agriculture, can induce antibiotic tolerance in *C. jejuni*. Importantly, antibiotic-tolerant *C. jejuni* cells act as a reservoir for developing FQ resistance because of elevated oxidative stress.

The development of FQ resistance occurred rapidly in *C. jejuni* during antibiotic tolerance. Among antioxidant enzymes available in *C. jejuni*, AhpC plays a major role in preventing the development of FQ resistance. Overall, the findings in this study provide novel insights into molecular mechanisms for antibiotic tolerance and resistance development and may explain the high prevalence of FQ resistance in *C. jejuni*.

## MATERIALS AND METHODS

### Bacterial strains and growth conditions

*C. jejuni* NCTC 11168 was used as WT in this study. The isogenic knockout mutants of Δ*ahpC* (43), Δ*katA* (44), Δ*sodB* (44), and Δ*perR* (45), and a CosR-overexpression strain (22) were reported in our previous studies, A *ΔperR* Δ*ahpC* double mutant and a CosR-overexpression strain with Δ*ahpC* were constructed by transforming the Δ*perR* mutant and the CosR-overexpression strain with the genomic DNA of the Δ*ahpC* mutant with natural transformation (46). *C. jejuni* strains were routinely grown at 42°C in Mueller-Hinton (MH) media (Difco) under microaerobic conditions (5% O2, 10% CO2, and 85% N2).

### Antibiotic tolerance assay

Overnight cultures of *C. jejuni* on MH agars were resuspended in 5 ml of MH broth in a 14 ml round bottom tube (BD Falcon, USA) to an optical density at 600 nm (OD600) of 0.08. The bacterial suspension was incubated with shaking at 200 rpm under microaerobic conditions. After 7 h, antibiotic exposure was initiated by adding 10x MIC or 100X MIC of antibiotics (ciprofloxacin, tetracycline, erythromycin, gentamicin). The concentrations were determined based on the MIC of WT (i.e., *C. jejuni* NCTC 11168). The 100x MICs of ciprofloxacin, tetracycline, erythromycin, and gentamicin were 6.3 μg/ml, 3.1 μg/ml, 100 μg/ml, and 50 μg/ml, respectively. For sampling, 1.2 ml of *C. jejuni* cultures were harvested and washed with fresh MH media. After washing, *C. jejuni* cells were resuspended in 100 μl of MH broth for enumeration onto MH agar plates and MH agar plates supplemented with 1 μg/ml ciprofloxacin. Colonies growing on MH agar plates supplemented with ciprofloxacin were randomly picked up and subjected to a broth microdilution susceptibility test to confirm resistance (47).

### Hydroxyl radical measurement

*C. jejuni* was treated with high concentrations of antibiotics for 24 h as described above. *C. jejuni* cells were washed twice with PBS and concentrated 10-fold. The assay was conducted according to the manufacturer’s instructions (Molecular Probes HPF, Thermo Scientific, USA). Briefly, 100 μl of a sample was placed on a 96-well plate (black opaque, Corning, USA). The hydroxyphenyl fluorescein (HPF) solution was diluted to the final concentration of 5 μM. Fluorescence at ex/em 530/590 nm was measured with a plate reader (Varioskan Flash; ThermoFisher Scientific) with gentle shaking at 25°C. Measured fluorescence signals were normalized to protein concentrations determined with a Bradford assay.

### Hydrogen peroxide measurement

The level of hydrogen peroxide formation under antibiotic treatment was measured with the Amplex Red Hydrogen Peroxide/Peroxidase Assay Kit (Invitrogen, ThermoFisher Scientific) according to the manufacturer’s protocol. *C. jejuni* was exposed to antibiotics for 24 h as described above. The samples were washed twice, concentrated 10-fold, and placed into a 96-well plate (black opaque, Corning). The working solution (10 mM Amplex Red reagent, 10 U/mL HPR stock solution, and reaction buffer) was added to each sample solution. The mixture was incubated at room temperature for 30 min, and fluorescence was detected at ex/em 530/590 nm with a plate reader (Varioskan Flash; ThermoFisher Scientific). Hydrogen peroxide concentration was determined by comparing with a standard curve prepared with known hydrogen peroxide concentrations and normalized to protein concentrations determine with a Bradford assay.

## ACKNOWLEDGMENTS

B.J. conceptualized the study. M.P., J.K., and J.F performed the experiments. B.J. and S.R. supervised the experiments. B.J., M.P., and K.S.L analyzed the results. B.J. wrote the manuscript. M.P. prepared figures.

This study was supported by funding from MnDRIVE (Minnesota’s Discovery, Research, and InnoVation Economy) to B.J and the Basic Science Research Program through the National Research Foundation of Korea (NRF) from the Ministry of Education (NRF-2021R1I1A1A01050990) to J.K.

We declare no competing interest.

