## Supplemental figures for "Development of fluoroquinolone resistance through antibiotic tolerance in *Campylobacter jejuni*"

**SUPPLEMENTAL DATA**

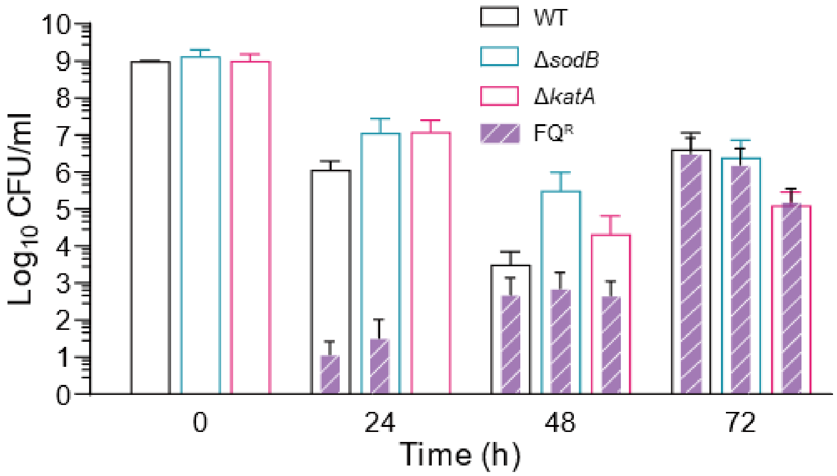


**Fig. S1. Fluoroquinolone (FQ) resistance development in Δ*katA* and Δ*sodB* mutants in the presence of 100x MIC of ciprofloxacin.** The results show the means and standard deviations of the levels of total *C. jejuni* (empty bars) and $FQ^R$ *C. jejuni* (patterned fills) of the results from three independent experiments.

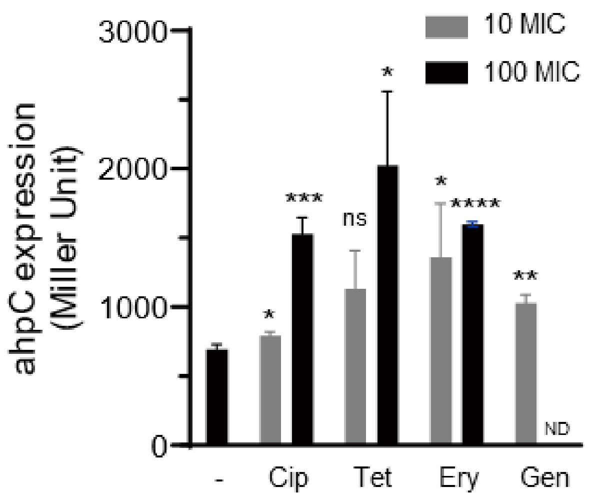


**Fig. S2. Increased transcriptional level of *ahpC* during exposure to high concentrations of antibiotics.** *C. jejuni* was exposed to 10x or 100x MIC of antibiotics for 24 hours before β-galactosidase assays. The assay was conducted as described previously (1). The results show the means and standard deviations of one representative experiment with three samples. The assay was repeated three times and produced similar results. Statistical analysis was conducted with Student's *t*-test. ns: non-significant, *: $P<0.05$, **: $P<0.01$, ***: $P<0.001$, and ****: $P<0.0001$.
